# Effects of SSRIs on the spatial transcriptome of dorsal raphe serotonin neurons

**DOI:** 10.1101/2024.10.10.617388

**Authors:** C. Henningson, J. Mlost, I. Pollak Dorocic

## Abstract

The serotonin system is the main therapeutic target for selective serotonin reuptake inhibitors (SSRIs) in treating depression, yet the mechanism of action of SSRIs remains incompletely understood. To investigate the molecular and transcriptional effects of SSRI administration on serotonin neurons, we performed spatial transcriptomics, a spatially resolved RNA-sequencing method in intact brain tissue. Mouse brain sections containing the dorsal raphe nucleus and adjacent midbrain structures were analyzed, revealing six distinct serotonergic subpopulations with unique molecular signatures and spatial distributions. Both acute and chronic fluoxetine treatment induced a large number of changes in gene expression in the dorsal raphe nucleus. Notably, *Htr1a* expression increased following acute treatment but decreased after chronic administration, supporting previous findings on serotonin transporter blockade effects on 5HT1a autoreceptors. Gene enrichment and network analysis identified key pathways modulated by SSRI administration, including Ras, MAPK and cAMP signalling pathways as well as pathways involved in axonal guidance. Additionally, we observed treatment-dependent opposing transcriptional changes in neuropeptides, particularly thyrotropin-releasing hormone (*Trh*) and Prodynorphin (*Pdyn*), with distinct spatial localization within the dorsal raphe nucleus. Collectively, our transcriptomic and in situ hybridization analyses reveal spatial and cell-type-specific heterogeneity in SSRI action within the dorsal raphe nucleus, providing new insights into the molecular basis of SSRI treatment effects.

## Introduction

Depression is a leading contributor to the global disability burden, with high lifetime prevalence, elevated risk of suicide, and a critically increasing incidence^1^. The serotonin system in the brain is the primary pharmacological target of the most common antidepressant treatment, the Selective Serotonin Reuptake Inhibitors (SSRIs). These drugs are routinely prescribed as a first-line treatment for depression and a range of comorbid conditions including anxiety and bipolar disorder, and are thus one of the most prescribed pharmaceutical drugs in the world. A large number of patients, in some countries as many as 10-15% of the total population^2^, currently rely on antidepressants to alleviate their suffering, with SSRIs being the most common form of treatment^3^. Despite this extensive clinical use, the molecular mechanisms underlying SSRI action remain incompletely understood. Furthermore, SSRIs exhibit variable treatment outcomes, delayed onset of therapeutic effects, and numerous side effects^4^. Understanding the mechanism of action represents a critical challenge in modern psychopharmacology and is essential for developing more effective therapeutic strategies.

Serotonin (5-hydroxytryptamine) is one of the major neuromodulators in the central nervous system, and plays a role in both normal function and disease. Serotonin interacts with at least 14 different receptor subtypes and therefore influences a wide range of physiological processes, from homeostatic maintenance to complex roles in mood and motivation^5^. In the mammalian brain, serotonergic neurons are exclusively located within the brainstem raphe nuclei, with the dorsal raphe nucleus (DR) containing the majority of forebrain-projecting serotonin neurons. These neurons are embedded within an extensive neural network, receiving inputs from multiple forebrain regions^6–8^ and projecting widely throughout the brain to modulate diverse behaviors^9^. One limitation of our current knowledge of the serotonin system, as well as its pharmacological targeting, is that the system has been considered unitary, *ie.* comprising all neurons that synthesize serotonin. However, recent single-cell RNA sequencing studies have described the heterogeneity of the serotonin system and elucidated a number of molecular serotonergic subpopulations within the DR^10–12^. The DR cytoarchitecture is arranged in a broad-level topography, with neuron types spatially arranged in the dorsal, ventral and later DR sub-compartments and linked to distinct forebrain-circuits. This cellular complexity is further enhanced by the presence of other neuronal types, including dopaminergic, glutamatergic, and GABAergic neurons, as well as various neuropeptide-expressing cells^13,14^. Understanding the spatial organization and molecular signatures of these distinct neuronal populations is crucial for elucidating their specific functional roles and responses to pharmacological interventions.

SSRIs exert their action by blocking the serotonin transporter (SERT), leading to rapid increases in extracellular serotonin concentration within hours^15^. However, clinical efficacy is only achieved after several weeks of chronic administration^16^. This delay in therapeutic effect is hypothesized to be related to 5-HT_1A_ receptor desensitization. 5-HT_1A_ are Gi-coupled receptors expressed both post- and pre-synaptically, with pre-synaptic autoreceptors providing crucial feedback inhibition of serotonin neurons. The initial SSRI-induced elevation of synaptic serotonin activates these autoreceptors, resulting in decreased serotonin neuron firing and release^17,18^. Chronic SSRI treatment leads to 5-HT_1A_ autoreceptor desensitization and subsequent loss of feedback inhibition, ultimately increasing serotonin neuron activity^19–21^. While this model has been influential in explaining the delayed therapeutic onset of SSRIs, alternative mechanisms have also been proposed, including enhanced hippocampal neurogenesis and increased synaptic plasticity^23^.

Antidepressants induce widespread transcriptional changes across multiple brain regions, including the prefrontal cortex, hippocampus, and amygdala^24–26^. However, most studies have focused on limited candidate genes in specific regions. Recent unbiased profiling using bulk RNA sequencing revealed that chronic fluoxetine treatment produces region-specific transcriptional effects across the brain, with the most pronounced changes occurring in the DR^27^. However, the specific effects on serotonin neurons, their spatial subtypes, and other cell types in the DR are unknown.

Here, we employed spatial transcriptomics and in-situ hybridization to analyze the molecular architecture of the mouse dorsal raphe nucleus under baseline conditions and following acute or chronic fluoxetine administration. We first established a high-resolution spatial map of the DR transcriptional landscape, identifying molecular markers that define distinct serotonergic subpopulations. We then characterized the transcriptional responses to SSRI treatment across these neuronal populations, revealing divergent effects of acute versus chronic administration. Our findings provide insight into the spatial and cell type-specific heterogeneity of SSRI action within the DR, with implications for understanding the complex mechanisms underlying antidepressant action. The data is accessible through an interactive app available at https://st-ssri.serve.scilifelab.se/app/st-ssri.

## Results

### Spatial transcriptomic profiling of midbrain and DR

Using unbiased and untargeted spatial transcriptomics, we profiled gene expression in coronal brain sections containing the DR and neighboring regions, in adult mice (**Fig. 1A**). We detected, on average, 3892 genes and 10,239 reads per spot (3892 ± 1133 unique genes, 10,239 ± 4978 reads). The complete dataset after quality control contained information on the expression of 30,413 unique genes across 8369 spots. Shared Nearest Neighbor (SNN) clustering of gene expression across the whole brain section revealed 13 distinct clusters (**Fig. 1B**). We did not observe batch nor sex mediated effects on clustering (**Fig. S1**). Aligning the molecular clusters with spatial location in brain tissue largely recapitulated anatomical brain regions, but also provided a higher-resolution delineation within the same brain region (**Fig. 1C**). For example, the periaqueductal gray (PAG) can be subdivided into the dorsal, lateral, ventrolateral and aqueductal PAG subregions with distinct molecular gene expression profiles. Spatial sequencing also clearly reveals the DR anatomical cluster, as well as a cluster flanking the DR bilaterally with distinct molecular signatures (**Fig. 1D**).

**Figure 1.**
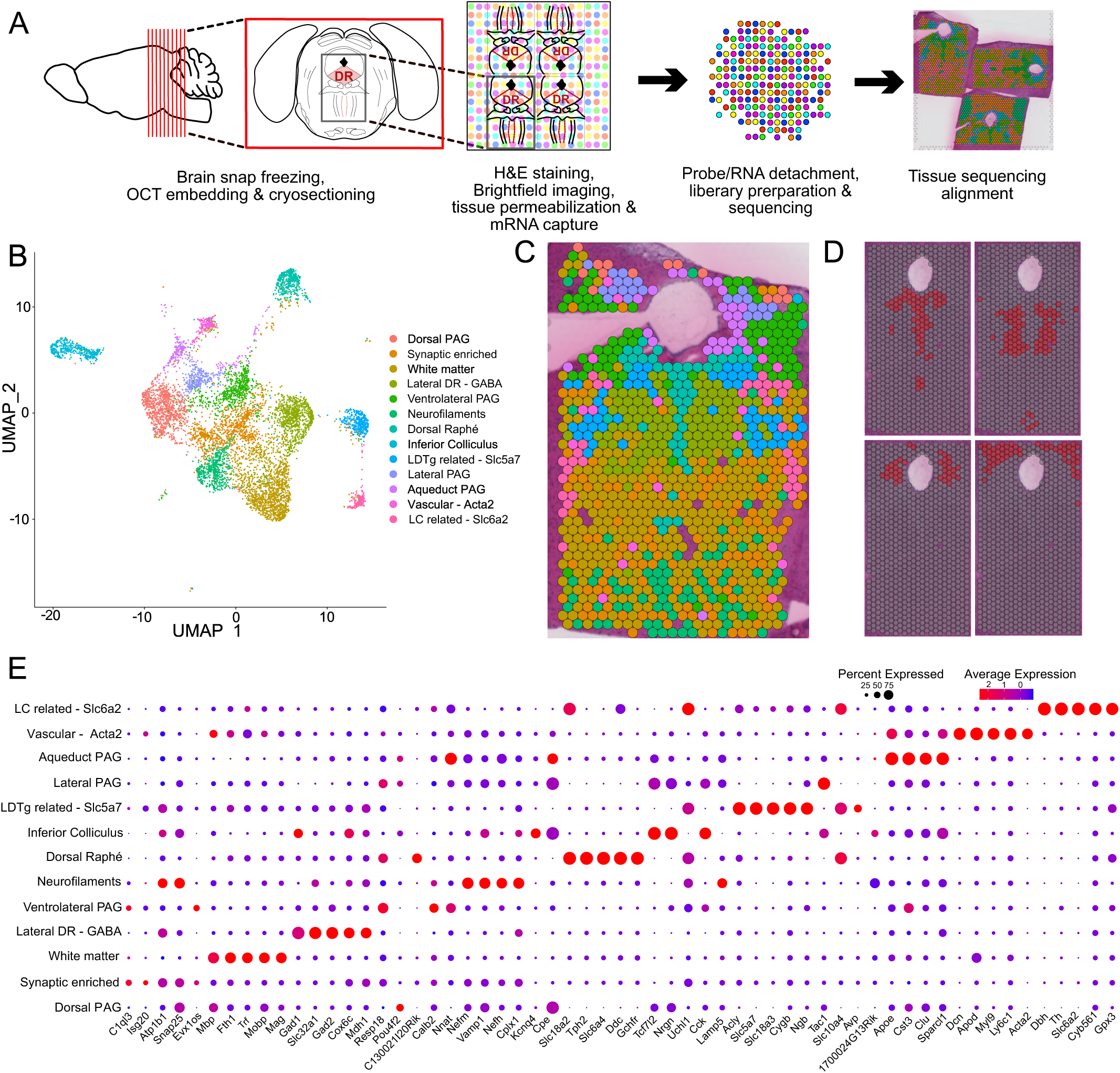
Spatial transcriptomic profiling reveals distinct molecular domains in the mouse midbrain and DR. **A.** Experimental strategy: Coronal sections of mouse midbrain including the DR were cut and multiple sections were mounted on Visium slide capture areas, followed by RNA sequencing and tissue alignment. **B.** UMAP plot based on the transcriptional signature of each spot, covering the whole midbrain section, revealed 13 molecular clusters. Acronyms in cluster names: PAG, periaqueductal gray; DR, dorsal raphe; LDTg, laterodorsal tegmental nucleus; LC, locus coeruleus. **C.** Representative midbrain tissue section with overlaid sequencing spots, color represents molecular cluster in (B). **D.** Visualization of four anatomically distinct spatio-molecular clusters, including DR (top left), lateral DR (top right), lateral PAG (LPAG, bottom left) and dorsal PAG (bottom right). **E.** Dot plot for each molecular cluster with expression level of individual genes. Diameter of the dot corresponds to the percentage of spots in the cluster expressing the gene, color indicates average expression of the gene.

The DR cluster is clearly identified with the expression of canonical serotonin markers, relating to serotonin synthesis, reuptake, release and regulation. These include: *Tph2* (Tryptophan hydroxylase 2), *Slc6a4* (serotonin transporter, SERT), *Slc18a2* (Vesicular monoamine transporter 2, VMAT2), *Ddc* (dopa decarboxylase), and *Gchfr* (GTP cyclohydrolase I feedback regulatory protein) that is involved in post-translational regulation of the serotonin synthesis pathway^28^ (**Fig. 1E**). Another molecular cluster (lateral DR-GABA), flanks the DR bilaterally and is marked by *Gad2* expression. Overall, the spatial sequencing approach revealed molecular markers for midbrain anatomical structures, and reliably identified the DR based on serotonin-associated gene expression. The subsequent analysis focuses on the DR cluster, in order to define spatial subpopulations of serotonin neurons, and determine the effects of acute and chronic fluoxetine treatment.

### Spatial subpopulations within DR serotonin system

The DR exhibits complex molecular heterogeneity across diverse cell populations. While single-cell RNA sequencing (scRNA-seq) studies have identified distinct serotonergic subpopulations^10–12^, this approach cannot preserve spatial information. Previous studies attempted to map the molecular subtypes to anatomical locations using visualization of limited candidate genes identified through scRNA-seq. However, no fully unbiased analysis of linking complete transcriptomic profiles directly to anatomical location in tissue has been performed in the DR.

Spatial transcriptomics analysis revealed 6 distinct molecular subclusters within the DR, each showing specific anatomical distributions along the rostro-caudal axis (**Fig. 2A,B**). These clusters demonstrated unique spatial organization: one midline, one ventral, one caudal, two lateral clusters, and one cluster with reduced serotonergic marker expression (**Fig. 2C**). All serotonergic clusters show strong molecular identity based on distinct marker genes, while the non-serotonergic cluster is defined by weak expression of canonical serotonin markers, likely containing less serotonin neurons compared to other DR spots. We identified cluster-specific marker genes, with several showing DR-restricted expression patterns: midline (*Slc17a8, Pdyn*), caudal (*Met*), and lateral cluster 1 (*Wfdc12, Trh*) (**Fig. 2D**). Other markers, including ventral cluster-associated *Apod* and lateral cluster 2-associated *Avp* and *Slc5a7*, showed broader expression patterns extending beyond the DR. Notably, lateral cluster 2 exhibited a distinct molecular profile, likely influenced by its proximity to cholinergic neurons of the rostral lateral dorsal tegmental nucleus, as supported by *Slc5a7* expression.

**Figure 2.**
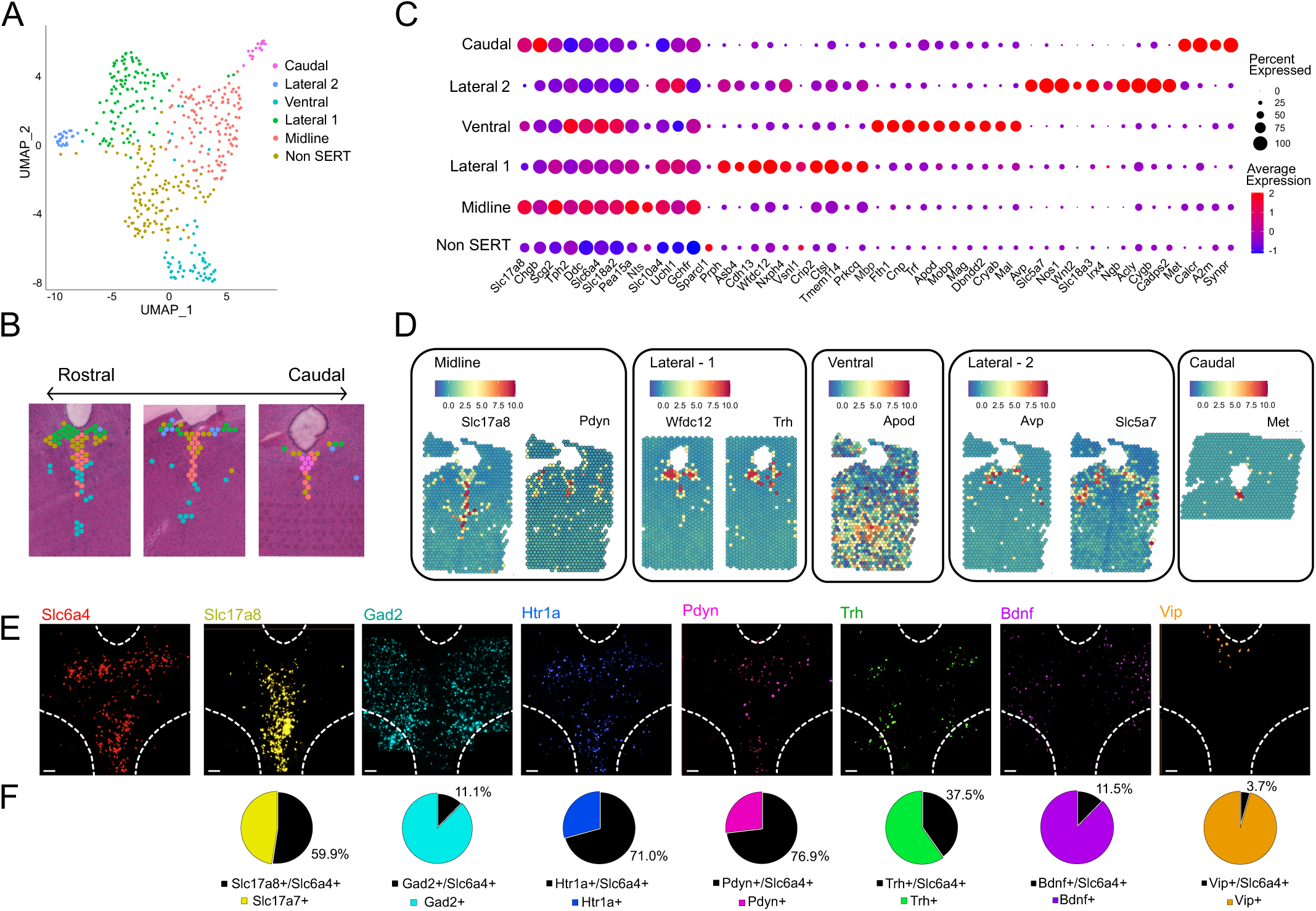
Spatio-molecular mapping of DR reveals specific spatial markers. **A.** UMAP plot based on the transcriptional signature of each DR spot, cluster names refer to the spatial location or molecular identity. **B.** Molecular clusters from A overlaid with tissue section, showing spatial localization of each cluster in DR and rosto-causal expression patterns. **C.** Dot plot showing expression of 10 cluster marker genes per cluster. Diameter of the dot corresponds to the percentage of spots in the cluster expressing the gene, color indicates average expression of the gene. **D.** Spatial heatmap of gene expression in DR for selected marker genes. Example of DR-specific markers (eg. midline-*Slc17a8*, *Pdyn*) and non-DR-specific marker genes (ventral-*Apod*). **E.** In-situ hybridization in DR (area within broken white border) with 8 probes*: Slc6a4, Gad2, Slc17a8, Htr1a, Pdyn, Trh, Bdnf, Vip,* each displaying a distinct single-cell distribution pattern within DR. Scale bar = 100µm. **F.** Quantification of the co-expression of each marker gene from (E) with serotonin marker *Slc6a4* (SERT).

To validate our spatial transcriptomics findings at single-cell resolution, we performed multiplexed in-situ hybridization (ISH) for key marker genes in the DR (**Fig. 2E, F**). The midline cluster is defined by the expression of *Slc17a8*, the gene coding for Vesicular Glutamate Transporter 3 (VGLUT3). This neural subpopulation has been extensively characterized, and has been shown to contain serotonin neurons that co-express VGLUT3 and are able to co-release glutamate in several forebrain regions^29–31^. We found that approximately half of *Slc17a8* neurons in the DR express *Slc6a4*, the gene coding for SERT (59.9% +/− 8.3% of *Slc17a8* cells express *Slc6a4*, mean ± SD, n = 1979 *Slc17a8* neurons, n = 6 mice) consistent with previous characterizations of this co-releasing population^13^. The GABAergic marker gene *Gad2,* which codes for Glutamate Decarboxylase 2, showed widespread expression throughout the DR and adjacent regions. A minority of DR *Gad2* cells co-expressed *Slc6a4* (11.1% +/− 5.1% of *Gad2* cells express *Slc6a4*, n = 3718 *Gad2* neurons, n = 5 mice). Analysis of the serotonin autoreceptor *Htr1a* revealed extensive but not exclusive co-expression with *Slc6a4* (71.0% ± 10.2% of *Htr1a* cells express *Slc6a4*, n = 3114 *Htr1a* neurons from 6 mice), confirming its predominant but not exclusive association with serotonergic neurons.

Our spatial transcriptomics analysis identified additional molecular markers with distinct spatial distributions in the DR. *Pdyn*, encoding the neuropeptide prodynorphin, showed strong midline expression and substantial co-localization with serotonergic neurons (76.9% +/− 6.5% of *Pdyn* cells express *Slc6a4*, n = 1235 *Pdyn* neurons, n = 6 mice). *Trh* (thyrotropin-releasing hormone) exhibited a specific expression pattern in the caudal and ventral lateral wings of the DR, with moderate serotonergic overlap (37.5% +/− 16.8% of *Trh* cells express *Slc6a4*, n = 483 *Trh* neurons, n = 5 mice). *Bdnf* (brain derived neurotrophic factor), linked to the regulation of stress and development of mood disorders^32^, showed predominantly lateral expression with limited serotonergic co-expression (11.5% +/− 6.9% of *Bdnf* cells express *Slc6a4*, n = 2709 *Bdnf* neurons, n = 5 mice). *Vip* (vasoactive intestinal peptide) was concentrated in the dorsal DR along the aqueduct, with minimal serotonergic overlap (3.7% +/− 3.0% of *Vip* cells express *Slc6a4*, n = 356 *Vip* neurons, n =6 mice). We integrated these findings into a comprehensive molecular map showing the spatial distribution and serotonergic co-expression patterns of each marker along the rostro-caudal axis (**Fig. S2**). These results reveal distinct molecular signatures that define spatially segregated neuronal populations within the DR.

Through integration of previously published single-cell RNA sequencing data from the DR^11^, we mapped five previously identified serotonergic subtypes (5-HT-I through V) onto our spatial transcriptomics dataset using CytoSPACE^33^ (**Fig. S3A**). CytoSPACE is a computational approach to integrate scRNA-seq and spatial transcriptomics data by assigning single cells to spatial locations based on gene expression similarity to determine cell type composition of each spot. Analysis revealed substantial overlap between certain subclusters and subpopulations, particularly evident between 5-HT-V and the Caudal subcluster (**Fig. S3A, B**) Analysis of top spatial cluster gene expression patterns demonstrated distinct molecular signatures across serotonergic subpopulations (**Fig. S3C**). The 5-HT-V subtype and Caudal spatial cluster exhibited concordant expression profiles, characterized by expression of *Met*, *Calcr*, and several other genes. The 5-HT-IV subtype aligned with the Midline spatial cluster, sharing *Slc17a8* expression while showing minimal expression of Lateral 1 marker genes. Similarly, 5-HT-I subtype and the Lateral 1 spatial cluster demonstrated shared expression of multiple markers, including *Prph*, *Cdh13*, and *Ctsl*. In contrast, subpopulations 5-HT-II and III displayed mixed expression patterns, incorporating markers characteristic of both midline and lateral populations (**Fig. S3C, D**). Quantitative spatial distribution analysis revealed preferential localization patterns, with 5-HT-II predominantly present in Lateral 1 (56% ± 13.3% of mapped 5-HT-II cells, mean ± SD, n = 37 of 66 neurons) and 5-HT-III showing enrichment in Midline regions (71% ± 4.5% of mapped 5-HT-III cells, mean ± SD, n = 17 of 24 neurons). These findings demonstrate that while serotonergic subtypes exhibit clear spatial preferences, each DR spatial cluster represents a distinct microenvironment characterized by specific combinations of 5-HT subtypes (**Fig. S5E**).

### Effects of acute and chronic SSRI treatment on DR serotonin system

SSRIs directly target serotonergic neurons by blocking SERT, yet their effects on gene expression within these neurons are not known. Previous transcriptional studies have focused primarily on downstream brain regions innervated by serotonergic projections, leaving a critical gap in our understanding of SSRI action at its primary site of effect. To address this, we employed spatial transcriptomics to analyze gene expression changes in the DR following fluoxetine administration. Given the well-documented temporal disparity between SSRI administration and therapeutic efficacy, we examined both acute and chronic treatment conditions to capture the dynamic transcriptional landscape underlying this therapeutic delay.

Three treatment protocols using gavage administration were administered to healthy, naïve mice: acute (single dose fluoxetine, 10mg/kg), chronic (22 days daily administration, 10mg/kg), and control (single dose of saline) (n=3/treatment group) (**Fig. 3A**). We sought to administer a dose and treatment schedule comparable to clinical use in treatment of depression^34,35^. Our analysis identified a range of differentially expressed genes (DEGs) between the control, acute and chronic treatment protocols in both the whole midbrain tissue clusters (**Fig. S4A)** and the DR clusters (**Fig. S4B,C**). Focusing specifically within the DR, acute and chronic SSRI administration resulted in DEGs enriched in several important signalling pathways and biological processes, including: Ras, MAPK, cAMP and axon guidance pathways (**Fig. 3B**). Chronic treatment led to widespread downregulation of axon guidance genes, most notably *Plxnc1*, which encodes a member of the plexin family and regulates axon guidance, cell motility and migration^36^ (**Fig. 3C**). Conversely, chronic treatment upregulated *Rasgrp1*, an activator of the Erk/MAP kinase cascade. The serotonin autoreceptor gene *Htr1a* showed biphasic regulation, with upregulation following acute treatment and subsequent downregulation after chronic administration, consistent with established SSRI-induced autoreceptor desensitization^19–21^. Similarly, the immediate early transcription factor *Fos* used as a proxy for neural activity, is downregulated over chronic treatment (**Fig. 3C**). These findings reveal broad temporal dynamics in gene expression across multiple functional pathways during SSRI treatment (**Fig. 3D, Fig. S4C**).

**Figure 3.**
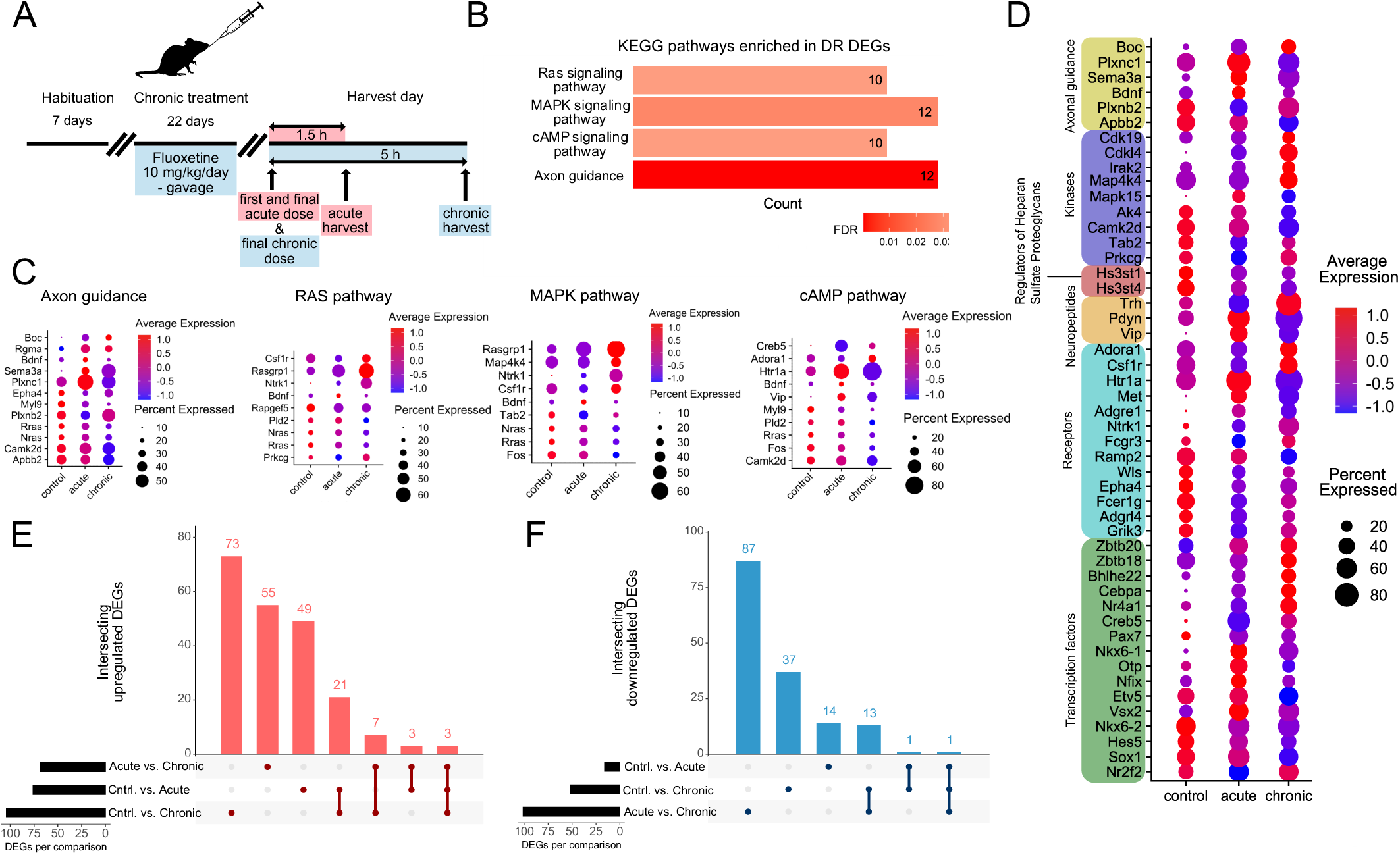
Acute and chronic SSRI treatment elicits distinct gene expression changes in the DR. **A.** Schematic representation of the pharmacological experiment. **B.** Gene enrichment analysis in the KEGG database of DEGs in the DR. Number in the bar represents the number DEGs observed across all treatment conditions. **C.** Expression profiles of single DEGs in the DR, categorized by the involvement in specific KEGG pathways. **D.** List of DEGs in the DR categorized by function from the Mouse Genome Informatics database and their expression levels across control, acute and chronic treatment conditions. **E.** Upset plot of upregulated genes in the DR, between treatment conditions. **F.** Upset plot of downregulated genes in the DR, between treatment conditions.

We investigated how many of the DEGs are uniquely modulated in each treatment condition by analyzing the degree of overlap between the DEGs in control, acute and chronic fluoxetine administration condition (**Fig. 3E, F**). From a total of 232 DEGs identified, 192 showed downregulation and 153 showed upregulation across all different treatment conditions. Comparing the treatments, the number of downregulated DEGs in the acute vs chronic treatment is highest, while each treatment comparison shows a large number of unique upregulated DEGs. Interestingly, very few up- and down-regulated DEGs are shared between all three treatment conditions.

In order to find key factors among DEGs, we performed network analysis of protein-protein interactions and selected the top three up- or down-regulated genes (**Fig. 4**). Following acute SSRI treatment, *Bdnf* was the only hub gene detected but among the top three upregulated genes were *Bdnf, Pdyn* and *Rgs5*, whereas the top three downregulated genes were *Myl9, Nptx2* and *Adgrl4* (**Fig. 4A**). Following chronic SSRI treatment, we found 7 hub genes: *Nras, Fos, Calb1, Rras, Fcer1g, Pdyn* and *Csf1r* (**Fig. 4B**). Among the top three upregulated genes were *Med1, Fcer1g, Csf1r*, whereas top three downregulated genes were *Pdyn, Nptx2* and *Fxyd5* (**Fig. 4B**). When comparing acute vs chronic SSRI treatment, we found 10 hub genes: *Fos, Met, Kit, Hcrt, Rbfox3, Trh, Ntrk1, Vip, Pdyn, Nkx2-2* (**Fig. 4C**). Among the top three upregulated genes were *Socs4, Bmp1* and *Bhlhe22*, whereas top three downregulated genes were *Vip, Tnnt1* and *Nkx6-1* (**Fig. 4C**). Interestingly, the two neuropeptide genes, *Pdyn* and *Trh*, showed opposite dynamic regulation over the time course of SSRI treatment. *Pdyn* was upregulated after acute and downregulated after chronic treatment, compared to the baseline. By contrast, *Trh* was initially downregulated, and then upregulated after the course of treatment (**Fig. 3D**).

**Figure 4.**
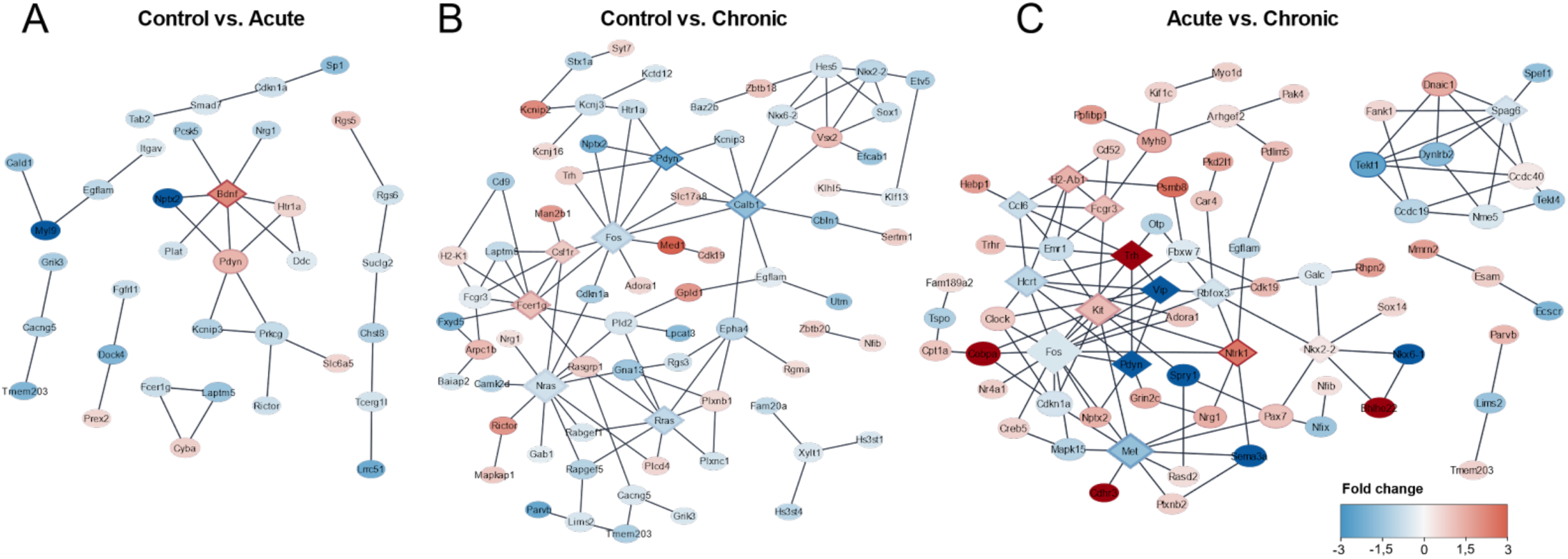
Network analysis of DR DEGs identifies key regulatory hubs across SSRI treatment. Protein-protein interaction network based on DEGs in DR in STRING database for changes in each comparison condition: control vs. acute (A), control vs. chronic (B) and acute vs. chronic (C). Hub genes are marked in a diamond node (◆). Red color denotes upregulation, blue denotes downregulation in comparison to control (A-B) or acute (C) group.

Analysis of neuronal activity patterns revealed distinct temporal responses to SSRI treatment across DR subregions (**Fig. S3D**). Using NEUROeSTIMator^37^, a bioinformatic approach that generates normalized activity scores based on the expression of 22 immediate early genes, we observed region-specific alterations in neuronal activity. Chronic SSRI administration resulted in significantly reduced overall activity in the DR compared to control conditions. This reduction in activity was specifically localized to the midline subcluster, while the lateral subcluster maintained baseline activity levels. Notably, this activity differential was only observed when comparing acute versus chronic treatment conditions. Analysis of the aggregate dataset showed no significant treatment effect on overall neuronal activity.

In summary, temporal analysis of SSRI administration revealed extensive transcriptional remodeling within the DR, with the most pronounced differences emerging between acute and chronic treatment conditions. These changes encompassed individual genes, molecular pathways, and region-specific activity patterns across DR subdomains. The transition from acute to chronic fluoxetine treatment was characterized by the largest number of differentially expressed genes, affecting both up- and down-regulated transcriptional programs and associated signaling networks.

### Dynamic gene expression changes of DR neuropeptides in acute and chronic SSRI treatment

Transcriptomic analysis revealed opposing regulation of *Pdyn* and *Trh* during SSRI treatment. To confirm these findings, we performed ISH on DR tissue after either acute or chronic fluoxetine treatment as well as with vehicle control. We found that acute administration increased both the number of *Pdyn*-expressing cells and total *Pdyn* mRNA spots in the midline DR (**Fig. 5A**). This acute upregulation was partially reversed following chronic treatment. Acute treatment also increased the number of neurons co-expressing *Pdyn* and *Slc6a4*. ISH analysis of *Trh* expression also corroborated our transcriptomic findings: chronic treatment increased both *Trh*-expressing cells and total *Trh* mRNA spots in the lateral DR (**Fig. 5B**). We observed no changes in *Trh* expression following acute treatment, nor in the number of neurons co-expressing *Trh* and *Slc6a4*. These findings demonstrate distinct regulatory mechanisms for *Pdyn* and *Trh*: SSRI-induced changes in *Pdyn* expression occur in serotonergic neurons, while *Trh* regulation appears independent of its co-expression with serotonin.

**Figure 5.**
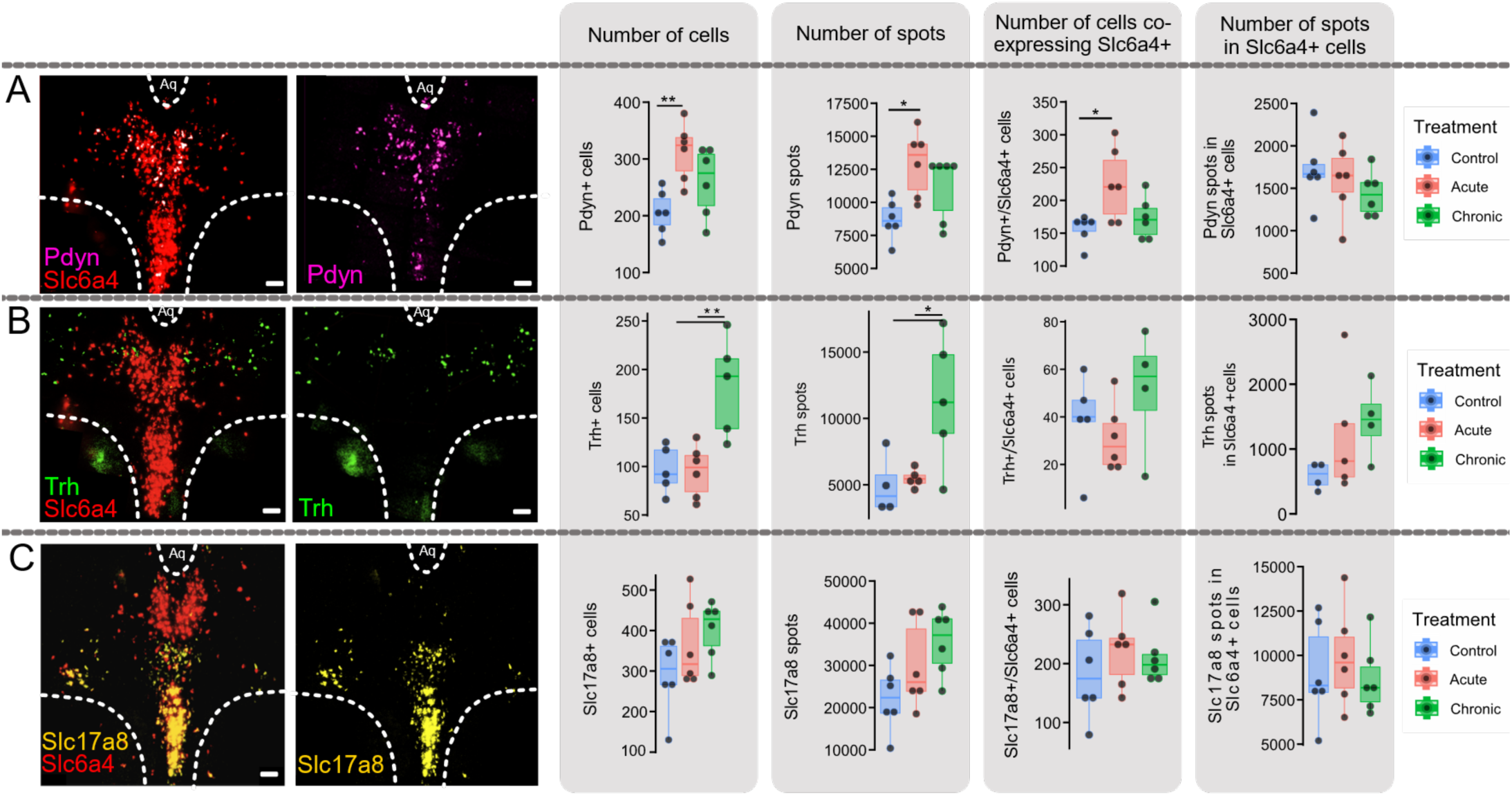
Dynamic gene expression changes of *Pdyn*, *Trh* and *Slc17a8* in DR following acute and chronic SSRI treatment. **A.** Representative images of in-situ hybridization of *Pdyn* and *Slc6a4* co-expression and *Pdyn* in the DR. Box plots comparing treatment groups: sum of *Pdyn+* cells per section (one-way ANOVA, F(2,15) = 6.64, p = 0.009**. Pairwise t-test with Holms correction; Acute vs Control, p=0.0072**), sum of *Pdyn* spots (transcripts) per section (one-way ANOVA, F(2,15) = 5.735, p = 0.014*. Pairwise t-test with Holms correction; Acute vs Control, p=0.0125*), sum of cells co-expressing *Pdyn* and *Slc6a4* per section (one-way ANOVA, F(2,15) = 5.004, p = 0.022*. Pairwise t-test with Holms correction; Acute vs Control, p=0.0255*), and sum of *Pdyn* spots in *Slc6a4* cells. Control n = 6, acute n=6, chronic n=6 mice per group. **B.** Representative images of in-situ hybridization of *Trh* and *Slc6a4* co-expression and *Trh* in the DR. Box plots comparing treatment groups of sum of *Trh* cells per section (one-way ANOVA, F(2,13) = 10.29, p = 0.002**. Pairwise t-test with Holms correction; Acute vs Chronic, p = 0.00132**; Chronic vs Control, p=0.00418**, control n = 5, acute n=6, chronic n=5 mice per group), sum of *Trh* spots (transcripts) per slice (one-way ANOVA, F(2,11) = 5.79, p = 0.019*. Pairwise t-test with Holms correction; Acute vs Chronic, p = 0.039*; Chronic vs Control, p=0.039*, control n = 4, acute n=5, chronic n=5 mice per group), sum of cells co-expressing *Trh* and *Slc6a4* per section (control n = 5, acute n=6, chronic n=4 mice per group) and sum of *Trh* spots in *Slc6a4* cells (control n = 4, acute n=5, chronic n=4 mice per group). **C.** Representative images of in-situ hybridization of *Slc17a8* and *Slc6a4* co-expression and *Slc17a8* in the DR. Box plots comparing treatment groups of sum of *Slc17a8* cells per section, sum of *Slc17a8* spots (transcripts) per slice, sum of cells co-expressing *Slc17a8* and *Slc6a4* per and sum of *Slc17a8* spots in *Slc6a4* cells. Control n = 6, acute n=6, chronic n=6 mice per group. Dots represent individual animals, p-values shown follow p < 0.05 = *, p < 0.01 = **, p < 0.001 = ***. Scale bar = 100μm. Boxplots show the data distribution in each group.

The gene *Slc17a8* (VGLUT3) has been speculated to play a role in the treatment effect of SSRIs^38^. We sought to clarify whether the expression levels of *Slc17a8* alone, or in cells co-expressing *Slc6a4*, are affected by SSRI treatment and found no evidence of either (**Fig. 5C**). Though identified as DEGs in the transcriptomic analysis, ISH revealed no significant treatment effects upon *Gad2*, *Htr1a*, *Bdnf*, or *Vip* expression in the DR (**Fig. S5**). Additionally, we investigated potential sex-specific effects of acute and chronic SSRI administration. The treatment-dependent regulation of *Pdyn* and *Trh* expression was consistent across male and female brains, while other markers showed no treatment effects in either sex. Overall, ISH analysis revealed no statistically significant sex-specific differences in the expression of *Pdyn*, *Trh*, *Slc17a8*, *Gad2*, *Vip*, *Bdnf*, and *Htr1a* across control, acute, or chronic treatment conditions (**Fig. S6**).

## Discussion

Using unbiased spatial transcriptomics, we generated a comprehensive molecular map of the DR, identifying six distinct subpopulations with unique spatial distributions. Our findings extend previous single-cell RNA-sequencing studies^10–12^ by providing crucial spatial context to DR molecular heterogeneity. We identified several region-specific markers: *Slc17a8* and *Pdyn* in the midline, *Met* in the caudal region, and *Wfdc12* and *Trh* in the lateral DR. These markers may be valuable candidates for targeted genetic manipulation of specific DR subpopulations using transgenic or viral approaches. Additionally, our analysis of neuropeptide expression revealed distinct spatial patterns, with *Pdyn*, *Trh*, and *Vip* showing region-specific distributions and varying degrees of co-expression with serotonergic markers.

Further, we investigated how DR gene expression is modified by SSRI treatment. Comparison of acute and chronic treatment conditions revealed extensive transcriptional changes. Gene enrichment and network analysis identified key modulated pathways among the 232 differentially expressed genes in the DR, with 10-12 genes annotated to each of the Ras, MAPK, and cAMP signaling cascades, as well as axonal guidance pathways. SSRI-associated deficits in axonal guidance have been documented, particularly during development^39^, and chronic fluoxetine treatment reduces both the density and morphology of serotonergic projections to the PFC and hippocampus^40,41^. Our findings show that chronic treatment decreased expression of several axonal guidance genes, including *Plxnc1*. Network analysis identified multiple RAS pathway regulators as hub genes. Changes in these hub genes and other highly regulated genes including *Bdnf*, *Csf1r*, *Nras*, *Rras*, *Fos*, *Met*, *Kit*, *Myl9*, and *Ntrk1* align with the neurotrophic hypothesis of antidepressant action^42^.

Our analysis revealed significant changes in neuropeptide expression, potentially underlying direct behavioral effects of SSRIs. Of particular interest was the regulation of *Pdyn*, which encodes prodynorphin, an opioid peptide that activates kappa opioid receptors (KOR). While KOR activation produces analgesia, it also induces dysphoria and depressive-like behaviors, distinct from other opioid receptor subtypes^43^. *Pdyn* expression occurs in multiple limbic regions, including the amygdala, hippocampus, nucleus accumbens, and hypothalamus, and has recently been identified in a subpopulation of DR serotonin neurons^10–12^. Although KOR signaling has been implicated in stress-induced depressive symptoms through its interaction with the dopaminergic system, its relationship with serotonergic function in mood regulation remains poorly understood. Studies using the forced swim test have shown that KOR disruption reduces depression-like behavior^44^, supporting KOR signaling’s pro-depressive role. Our findings revealed biphasic regulation of *Pdyn*, with increased expression following acute SSRI treatment and subsequent decrease after chronic administration. This initial upregulation may contribute to the documented adverse effects during early SSRI treatment, including increased anxiety and suicide risk^45,46^.

The neuropeptide-coding gene *Trh* showed opposing temporal regulation compared to *Pdyn*, with downregulation after acute treatment and upregulation following chronic SSRI administration. While thyrotropin-releasing hormone (TRH) is primarily known for its role in the hypothalamic–pituitary–thyroid axis, it is also expressed in limbic regions, including the hippocampus, amygdala, and lateral DR. TRH signals through two receptors (TRHR1 and TRHR2) expressed throughout these regions, suggesting broader functions beyond endocrine regulation. Evidence supports TRH involvement in cognitive and emotional processing^47^, with both TRHR1 and TRHR2 knockout mice exhibiting depression- and anxiety-like behaviors^48,49^. Clinically, depression is associated with blunted TRH response and hypothyroidism^47^, though the distinct roles of region-specific TRH systems remain unclear. Our observation that chronic SSRI treatment upregulates *Trh* expression in the DR suggests a potential mechanism contributing to therapeutic efficacy. Notably, neither *Trh* nor *Pdyn* regulation showed sex-specific effects during SSRI treatment.

Our findings provide molecular support for the 5HT_1a_ desensitization hypothesis, with *Htr1a* showing upregulation following acute treatment and downregulation after chronic administration. We observed reduced *Fos* expression after chronic SSRI treatment, suggesting decreased neuronal activity. Chronic treatment particularly suppressed activity in the midline DR cluster. These transcriptional changes present an interesting contrast with previous electrophysiological and immunohistochemical studies, which showed acute SSRI treatment initially decreases serotonergic neuron activity before returning to baseline during chronic treatment, with variable region-dependent effects^17,50–52^. This apparent discrepancy might be explained in two ways. First, our spatial transcriptomics data represents mini-bulk measurements, potentially capturing SSRI-induced changes in non-serotonergic neurons within the analyzed regions. Alternatively, distinct serotonergic subtypes may respond differently to SSRI treatment, with some populations showing decreased activity while others maintain or increase their firing rates. Future cell type-specific studies will be crucial to resolve these possibilities and elucidate the temporal dynamics of SSRI effects across distinct neuronal populations.

Our analysis reveals spatial molecular heterogeneity within the DR and provides new insights into region-specific SSRI actions. While our findings support established theories of SSRI mechanisms, including 5HT_1a_ desensitization and neurotrophic effects, we also uncovered a potential novel neuropeptide-based mechanism involving dynamic regulation of *Pdyn* and *Trh*. The opposing temporal patterns of these neuropeptides may contribute to both the initial adverse effects and subsequent therapeutic benefits of SSRI treatment. The candidate genes, pathways and cell types uncovered pave the way for deeper mechanistic studies to unravel the underpinnings of depression, and reveal novel targets for more specific pharmacological targeting.

## Acknowledgements

This study was supported by SciLifeLab and Wenner Gren Foundations grants to I.P.D, and Polish National Agency for Academic Exchange (The Bekker Programme - BPN/BEK/2021/1/00152) and SciLifeLab RED Postdoctoral Fellowship to J.M. The authors acknowledge support from the National Genomics Infrastructure in Stockholm funded by Science for Life Laboratory, the Knut and Alice Wallenberg Foundation and the Swedish Research Council, and SNIC/Uppsala Multidisciplinary Center for Advanced Computational Science for assistance with massively parallel sequencing and access to the UPPMAX computational infrastructure. This work was supported by the National Bioinformatics Infrastructure Sweden (NBIS) at SciLifeLab. We thank A.R. Costa, and all the members of the Pollak Dorocic lab for thoughtful discussion, and C. Broberger for generous use of technical equipment.

## Author contributions

C.H. performed the experiments, analysed and visualised the data. J.M. contributed to spatial transcriptomic analysis and produced the network analysis. I.P.D. conceived and supervised the study. All authors wrote and edited the manuscript.

## Data availability

Data has been deposited at <u>10.5281/zenodo.14859692</u> and is also accessible through an interactive app available at https://st-ssri.serve.scilifelab.se/app/st-ssri. Any additional information and code details are available from the lead contacts upon request.

## Methods

### Animals

All experiments were carried out in adult C57BL/6J (Charles River) male and female mice. Mice were maintained under standard housing conditions with a 12-hour light cycle and with ad libitum access to food and water. All animal experiments had received approval from the local ethical board, Stockholms Norra Djurförsöksetiska Nämnd, and were performed in accordance with the European Communities Council Directive 2010/63. For spatial transcriptomics experiments, 9 mice (5 male, 4 female) were used, for *in-situ* hybridization 17 mice (9 females, 8 males) were used.

### Fluoxetine administration

Fluoxetine Hydrochloride (Sigma-Aldrich, F132) was dissolved in saline and stored at −20C° until use. For administration the fluoxetine hydrochloride was diluted to 2 mg/ml and administered to the mice by gavage in 10 mg/kg dose. The dose was calculated to be within the range of human equivalent doses prescribed for depression. The standard dose of fluoxetine for treatment of depression in humans is 20mg/day, with the highest being 80 mg/day^1^. Assuming an adult human weighs 60kg,the range of doses are 0.333-1.333 mg/kg. converting the 10 mg/kg mouse dose into the human equivalent is 0.81 mg/kg^2^.

The gavage procedure was performed with soft flexible feeding tubes to minimize the discomfort and potential damage to the animal. Mice in the acute treatment group received a single 10mg/kg dose of fluoxetine and brain tissue was harvested after 1,5h. Fluoxetine has been shown to produce behavioural effects in C57BL/6J mice after 30 min^3^. Mice in the chronic treatment group received 10 mg/kg/day for 22 consecutive days. The brain was harvested 5 h after the final dose to avoid any acute dose effect from the final dose. The control group received a single dose of saline instead at the equivalent volume to weight followed by brain harvest after 1.5 h.

### Tissue preparation

Isoflourane was used to render the animals unresponsive before cervical dislocation. The animals were decapitated, the brain extracted and immediately placed in ice cold saline for a maximum of 15 sec. For spatial transcriptomics experiments, the midbrain was dissected on an ice cold metal block of approximately 5×5×2 mm. The brains utilized for RNAscope experiments remained whole. The tissue was flash frozen using powdered dry ice and was stored airtight at −80C° prior to sectioning. All tools and surfaces were cleaned with 70% ethanol and RNAase away (Thermo Scientific, 10666421) prior to and during the harvesting.

For spatial transcriptomics, the dissected midbrains were trimmed, labelled and embedded together in O.C.T. (VWR, 361603E). The blocks were sectioned on a cryostat in 10 µm coronal sections that were placed on the Visium slide capture areas. 3-4 trimmed brain sections were placed in single capture areas. For RNAscope experiments, whole brains were embedded in O.C.T., cut in 20 µm coronal sections on the cryostat and placed on SuperFrost+ slides and allowed to dry in the cryostat cold chamber. The slides were stored airtight at −80C° until either RNAscope or Visium experiment start.

### Fluorescent in-situ hybridization

*In situ* hybridization to detect expression of genes of interest was performed using Hiplex RNAscope technology following the manufacturer’s protocol for fresh frozen tissue (Advanced Cell Diagnostics, ACD, Hayward, CA). In brief, 20µm sections were fixed in 4% PFA (Sigma-Aldrich, 252549) solution (60 min, RT), dehydrated for 5 min each in 50%, 75%, 99% and 99% laboratory grade ethanol (Fisher Chemical E/0600DF/F17). The slides were stored overnight in 99% ethanol at −20C°. The next day the slides dried at RT and a hydrophobic barrier was drawn around each tissue section. Protease IV was added to permeabilize the tissue sections (30 min, RT). After washing, each section was covered in probe dilution of 8 probes, or with positive and negative control probes, and incubated at 40C° for 2h in a humidity controlled environment. With wash steps between each incubation three amplification steps were performed, each 30 min 40C°. To reduce autofluorescence the slices were incubated with 3% FFPE reagent mix 30 min RT prior to ‘HiPlex Fluoro T1–T4 v2’, 15 min 40C°. After washing, DAPI staining was applied followed by clear mounting media and the first round of imaging.

The next day the sections were washed, and fresh cleaving solution was applied for 15 min RT two consecutive times, with washes in between to ensure removal of signal from previous imaging round. The slides were washed and incubated with 3% FFPE reagent mix 30 min RT, followed by ‘HiPlex Fluoro T5–T8 v2 ‘ for 15 min 40C°. After washing, clear mounting media was added, followed by the second imaging round. The following probes were used: Htr1a-T1, Vip-T2, Bdnf-T3, Pdyn-T4, Trh-T5, Slc6a4-T6, Slc17a8-T7, Gad2-T8.

### Microscopy

Imaging was performed on a Leica DM6 B stand up microscope with 40x oil immersion lens using the Leica developed LasX software. Intensities and exposure were setup on a separate tissue to minimize tissue bleaching. Each section was scanned in brightfield and the image ROI was selected and saved. Several focus points were added for each ROI to ensure uniform focus throughout the stitched image. Each slice was imaged in 5 channels (DAPI, 488, 550, 650, 750) with a z-stack of 8×8 um depth over two separate imaging rounds. The LasX Mosaic Merge stitching tool was used to merge the individual images together, using the blend ‘smooth’ setting.

### Image analysis

All images were imported to Fiji (ImageJ 2.14.0), using Bio-Formats importer to create a single max intensity projection image for all four channels. The max intensity image from imaging round 1 & 2 of the same slice was uploaded into a Qupath (0.5.0) project^4^. Using the Fiji/Qupath plugin, BigWarp in Fiji was utilized to create coordinate maps to overlap the two rounds of imaging based on the DAPI staining in the same slice.

Cell segmentation was performed in Qupath with Stardist scripts using their dsb2018_heavy_augment.pb model, identifying nuclei based in DAPI staining with 5.0 cell expansion and 0.5 threshold^5^. A region of interest was selected that included all Slc6a4+ cells, and corresponded with the Dorsal Raphe Nucleus location. Within this ROI Qupath’s own ‘Object Classification’ was utilized to assign cell identity based on marker expression. Object classifiers were trained either on the image itself or on other images in the data set. Individual RNA puncta were quantified using Qupaths subcellular detection feature. The detection threshold was set individually for the image based on a combination of the autofluorescence of the image and the fluorescence strength seen in the negative control for the specific channel.

Objects and annotations and their measurements were transferred to the Round 2 image using the Warpy plugin “transfer annotations and measurements”.

Two rounds of exports were merged based on the shared cell object ID into a single data frame saved per animal and slice. All slices were combined into a larger data frame from which summary tables were produced to assess sum or cells, sum of spots, both per marker and co-expressing subpopulations. We tested our data for normal distribution and initially performed ANOVA to compare our three treatment groups, followed by pairwise t-test adjusted with the Holm-correction, if the ANOVA was statistically significant (p<0.05).

### Spatial transcriptomics

The RNA integrity of the frozen tissue sections was measured with a bioanalyzer (Agilent) and was above 8 RIN for all samples. The 10x Genomics Visium protocol was performed by National Genomics Infrastructure at SciLifeLab, Sweden. Briefly, the sections were fixed with methanol at −20C° before performing H&E staining and bright field imaging. The tissue was then permeabilized enzymatically and the RNA was captured on the barcoded probes attached to the glass slide by the poly-A tail. The remaining tissue was washed away, and libraries were generated for each capture area and sequenced. Manual alignment of the H&E stained histology images to the expression spot grid was performed using the 10x Genomics Loupe Browser software (v. 5.1.0). The raw sequencing data files (FASTQ files) for the sequenced library samples were processed using the 10x Genomics Space Ranger software (v. 1.3.0) using the mouse genome reference transcriptome version GRCh38 2020-A (July 7, 2020) provided by 10x Genomics.

### Analysis of Visium sequencing data

For each capture area the respective histological image was used for data alignment. The data was processed in R 4.2.2 with Seurat (V4.3.0).

Spots were filtered based on cutoff values on several QC parameters: spots containing less than 1000 genes, spots containing less than 1000 nFeatures, spots containing more than 24 % mitochondrial genes. Genes from a number of categories were removed from further analysis, which included haemoglobin genes, mitochondrial genes, ribosomal genes and X/Y chromosome genes. For each capture area, data was normalized using SCTransform with ‘glmGamPoi’ method, v2 vst.flavor and regressing out % mitochondrial genes and % ribosomal genes. Only a set of variable genes was returned for further analysis. In order to minimize batch effects, data across the capture areas was integrated using Reciprocal PCA based on top 3000 features and top 30 principal components. To find integration anchors we used capture area containing slices from the control-treated group as a reference and K = 10.

Clustering was performed using the original Louvain algorithm and resolution = 0.9. To visualize data points in two-dimensional space, we have performed Uniform Manifold Approximation and Projection (UMAP) on the first 20 principal components using 50 closest neighbours and spread = 2. Three clusters were excluded based on criteria that it must be present on a minimum two slices per condition. After initial clustering, we selected a cluster with elevated Tph2 expression as a DR cluster and subjected it to subclustering by using the same algorithms but changed the number of dimensions to top 8 and resolution to 0.8. We have carefully selected above mentioned algorithms and parameters by comparing clustering stability, purity and silhouette width. To find markers and perform differential gene expression analysis, we used FindMarkers() function on “data” slot from integrated dataset and compared medians by Wilcoxon Rank Sum test. We have set “logfc” threshold to 0.5 and minimum expression in at least 20% of cluster’s spots. Differential gene expression was analysed comparing a single cluster with the rest of the dataset to find cluster marker genes and within the same cluster between treatment conditions to find treatment affected genes. For multiple comparison correction, we used False Discovery Rate and only changes with p-value < 0.05 were considered significant.

#### Network analysis

To identify key genes in DR affected by SSRI treatment, we created a Protein–Protein Interaction (PPI) Network with the STRING database using the list of all DEGs detected in DR, with significance cutoff value of FDR < 0.05 and minimum expression in at least 20% of the cluster spots. Network was then visualized and analyzed in Cytoscape (3.10.0). Based on the degree distribution in all three networks, the degree threshold value for hub genes was set to 6. The PPI network incorporates protein interactions known from experiments, text mining, databases, co-expression, co-occurrence, neighbourhood and gene fusion.

#### Gene enrichment analysis

In order to investigate the potential functional effects of the SSRI treatment in the DR we performed gene enrichment analysis through The Database for Annotation, Visualization and Integrated Discovery (DAVID)^6,7^ bioinformatics. We uploaded all DEGs detected in the DR, with significance cutoff value of FDR < 0.05 and minimum expression in at least 20% of the cluster spot, as a list in the DAVID analysis wizard and selected annotation categories KEGG pathways and GO terms Biological processes_5.

#### Activity analysis with NEUROeSTIMator

Using the gene-level counts per Visium spot as input for the NEUROeSTIMator^8^ v0.1. One-way Anova with following Tukey’s test were performed for statistical analysis.

#### Deconvolution with CytoSPACE, mapping single cell seq data to Visium

Huang et al, 2019 single cell sequencing data was used for cell type deconvolution performed with CytoSPACE^9^ v1.0.5, default settings. Estimated number of cells was set to 9 based on VistoSeg^10^. All available cell types were mapped, but only serotonin subtypes were isolated for visualization and further quantification.

### Reproducibility and sample size

Sample size based on other previous research with no calculations done prior to the experiments. All experiments contained between 3-6 biological replicates per group. Data was excluded based on raw data quality, if dissection of the slices did not contain the same clusters in all treatment groups, or if the cluster contained too few data points for further analysis.

---

**Figure S1.**
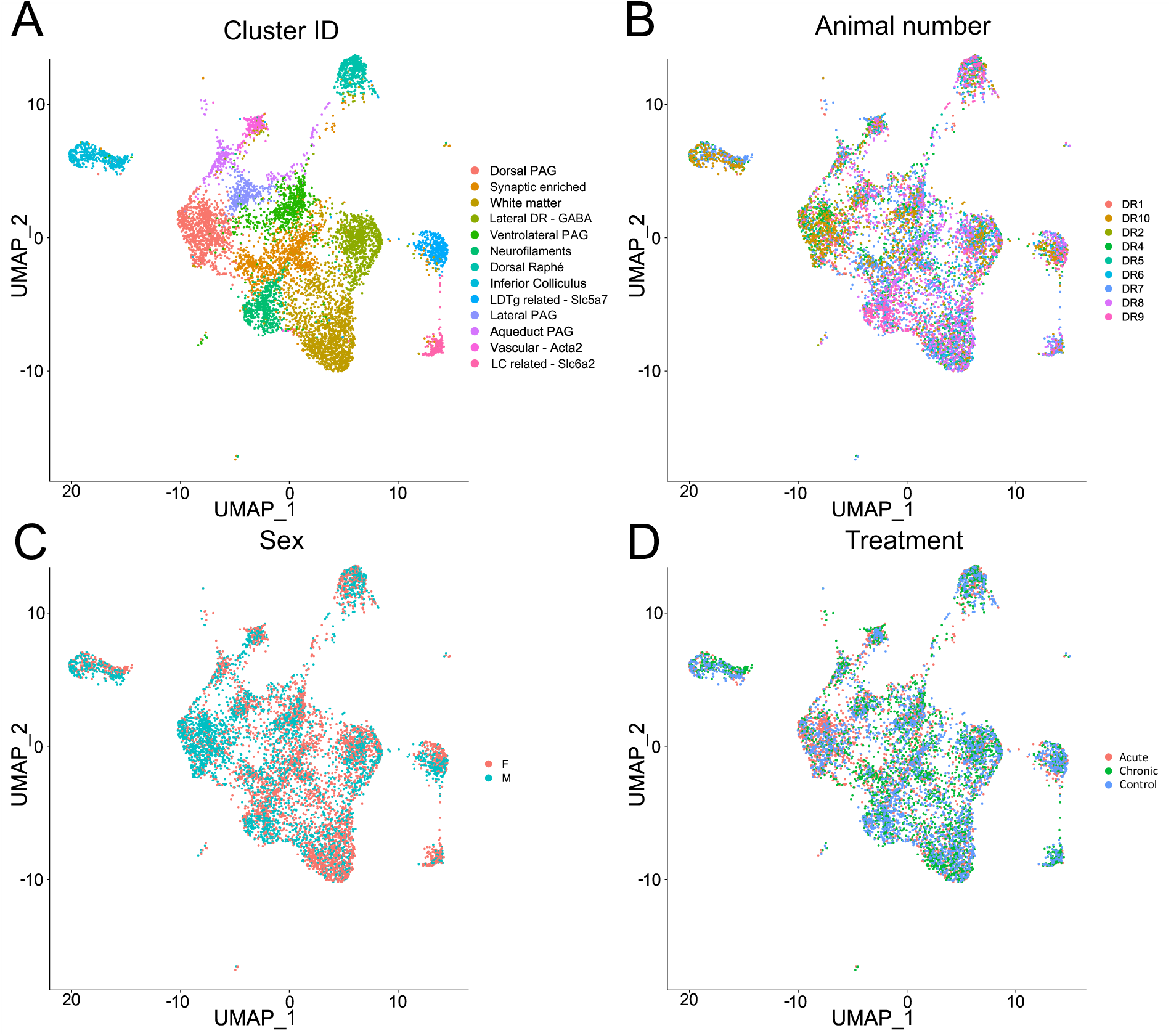
UMAP plot of midbrain transcriptomic data show no effect of sample, sex, and treatment in clustering. Lack of the technical- or batch-driven effects on clustering results in UMAP plot (**A**) was visualized by overlaying sample ID from different animals (**B**), their sex (**C**) or treatment group (**D**).

**Figure S2.**
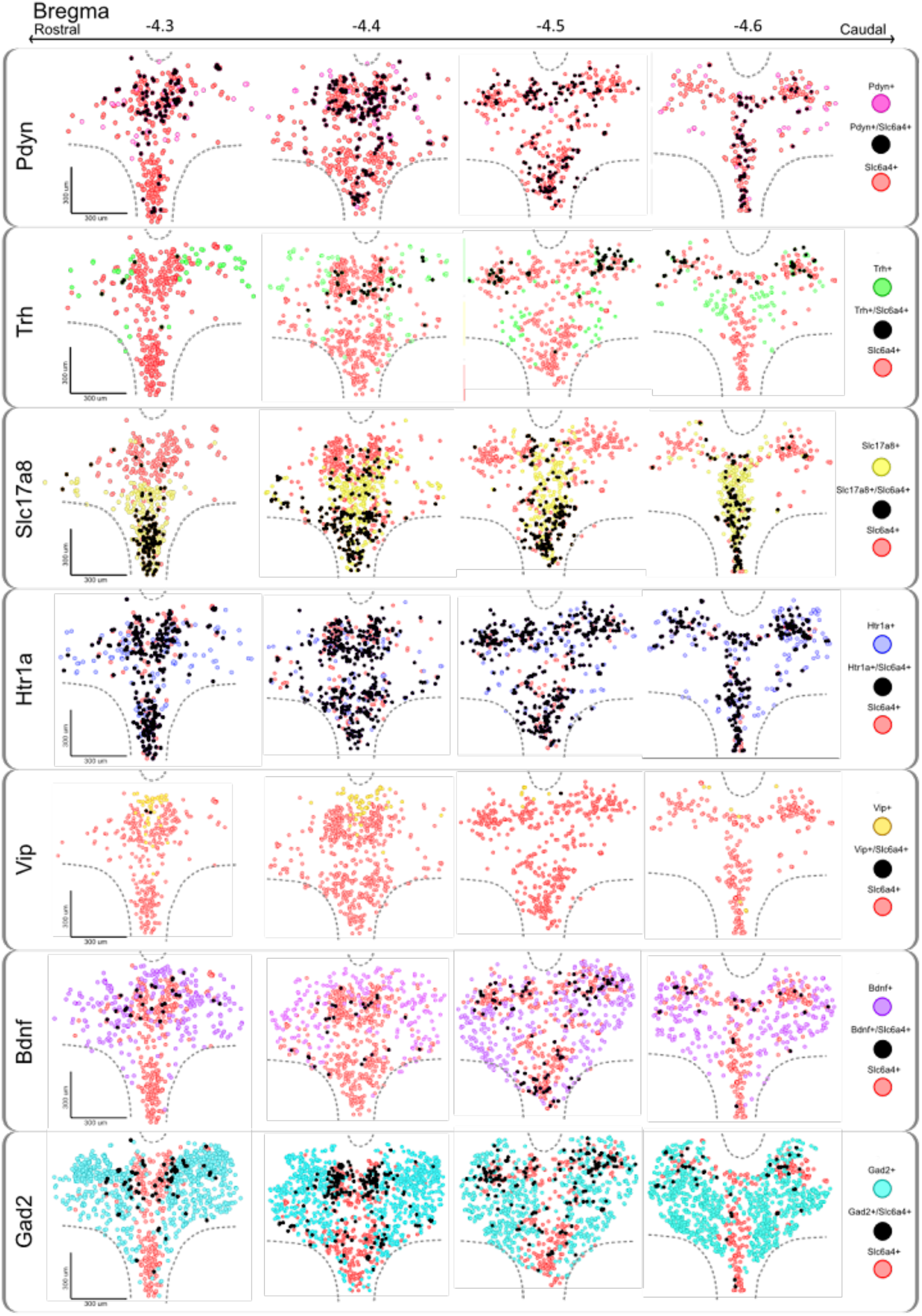
Regional expression and co-localization with serotonin for multiple markers in the DR along the rostro-caudal axis. Spatio-molecular map for each marker gene (*Pdyn, Trh, Slc17a8, Htr1a, Vip, Bdnf, Gad2*) in the DR, along the rostro-caudal axis (−4.3mm to −4.6mm). Co-expression of each specific marker with *Slc6a4* (SERT) is marked by a black circle (each circle correspond to a single cell), Slc6a4 positive cells lacking marker co-expression are marked in red, and the specific marker cells lacking Slc6a4 expression are shown in their respective color.

**Figure S3.**
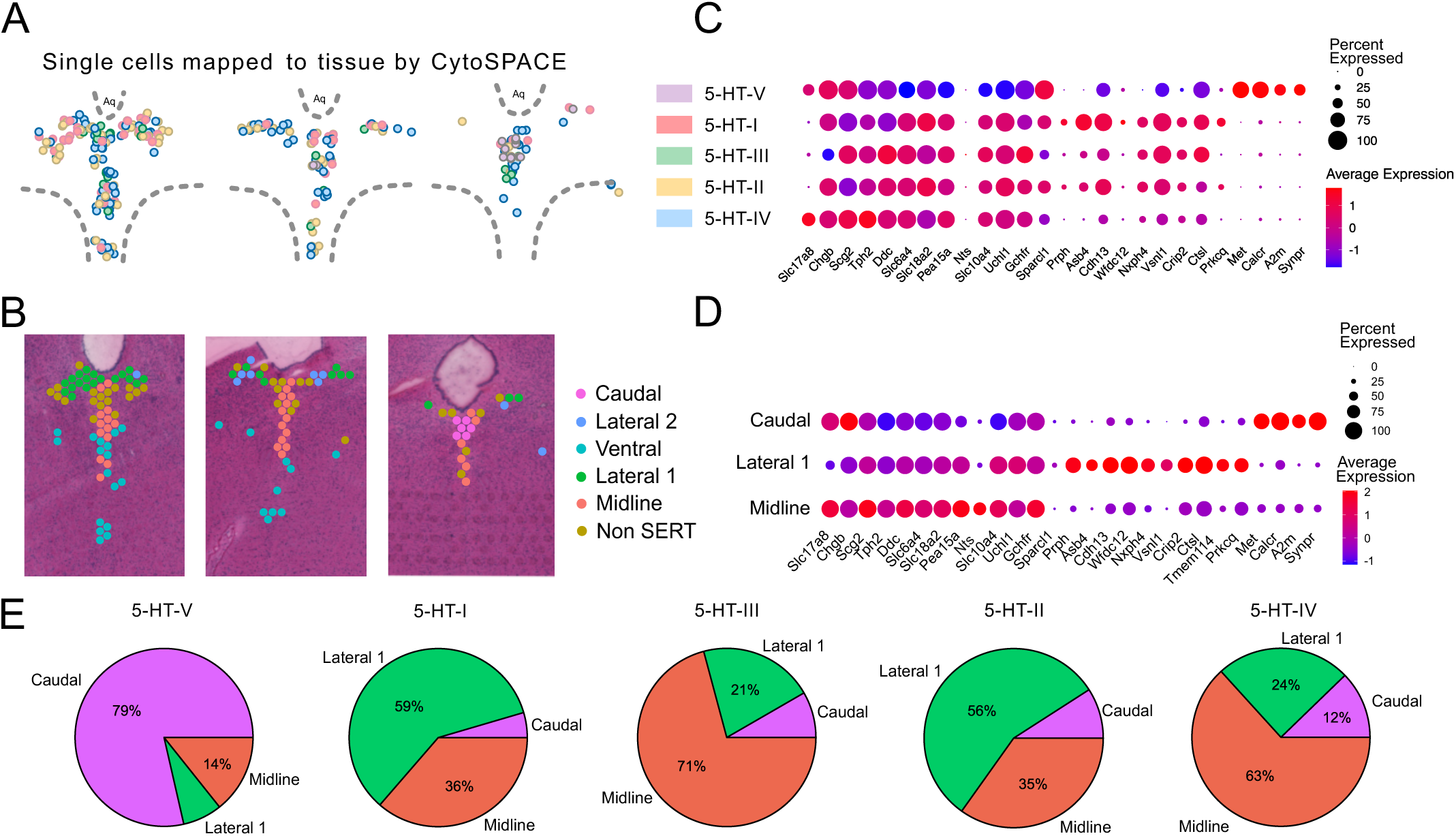
Mapping and deconvolution of serotonin subtypes reveals distinct spatial domains in DR. **A.** Integration of five molecular subtypes of DR serotonin neurons from published scRNA-seq dataset (5-HT-I through V from Huang, et al. 2019) mapped to spatial location across the rostro-caudal axis, using CytoSPACE computational deconvolution. **B.** Spatial transcriptomic clusters overlaid with tissue section, showing spatial localization of each cluster in DR and rostro-caudal expression patterns. **C.** Expression of top genes from spatial clusters (Caudal, Lateral 1 and Midline) mapped to five serotonin subtypes (5-HT-I through V). Diameter of the dot corresponds to the percentage of spots in the cluster expressing the gene, colour indicates average expression of the gene. **D.** Expression of top genes from spatial clusters (Caudal, Lateral 1 and Midline). Diameter of the dot corresponds to the percentage of spots in the cluster expressing the gene, colour indicates average expression of the gene. **E.** The distribution of serotonergic subtypes (5-HT-I through V) was quantified across three spatial domains (Caudal, Lateral 1, and Midline), expressed as the percentage of each subtype’s total mapped cells within these clusters.

**Figure S4.**
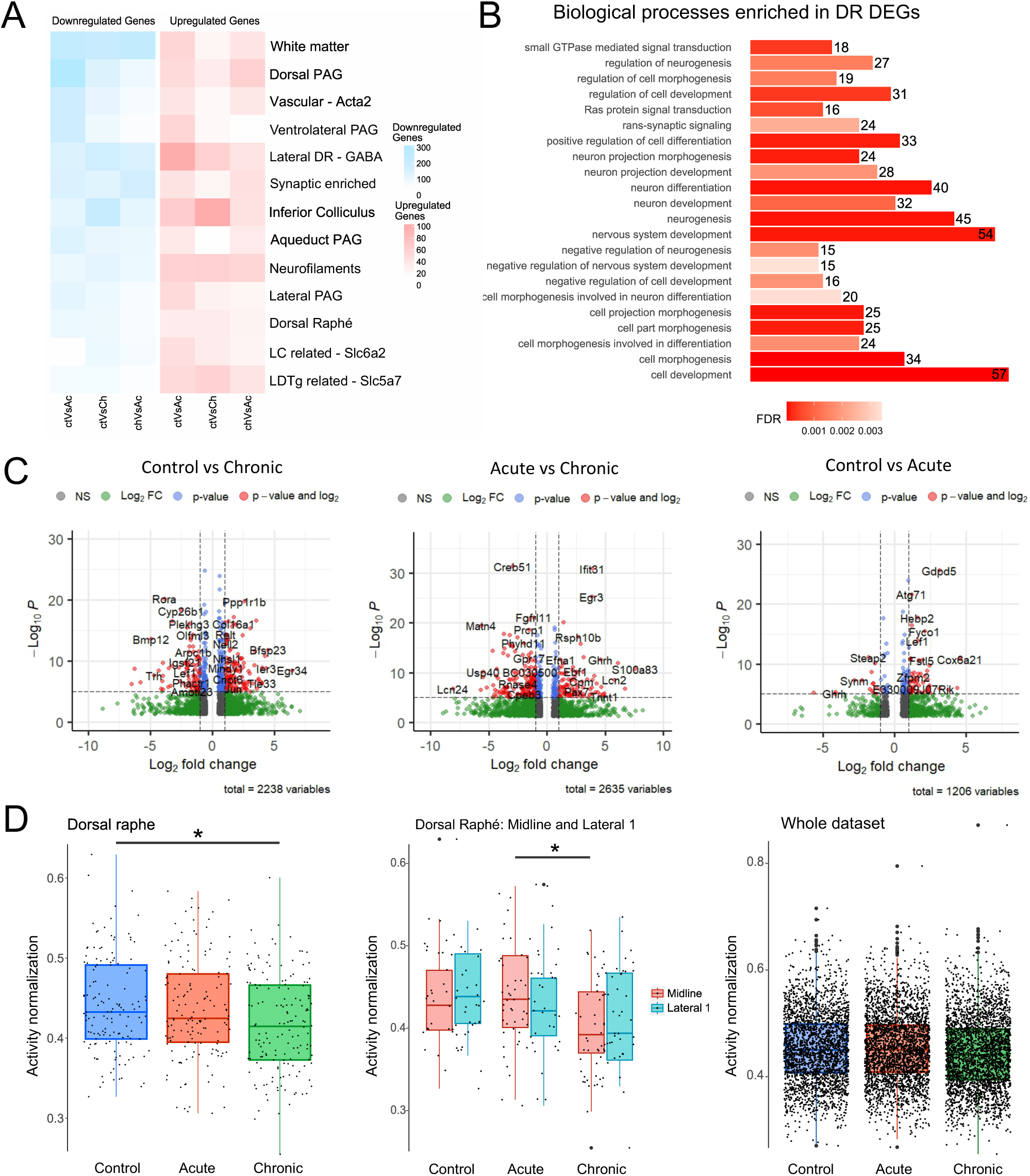
SSRI-induced changes in gene expression and activity in midbrain and DR. **A.** Heatmap visualization of the number of DEGs in clusters from whole midbrain section, across treatment conditions. The clusters correspond to UMAP in Fig. 1B. **B.** Results of gene enrichment analysis in Gene Ontology, Biological Processes of DEGs in the DR across all treatment conditions. **C.** Volcano plots representing top up- and down-regulated genes in the DR across treatment conditions. **D.** Box plot comparing DR cluster neural activity between treatment groups, activity per transcriptomic spot generated by NEUROeSTIMator (one-way ANOVA followed by Tukey multiple comparisons of means, F(2,443) =4.101, p =0.0172*; Control vs Chronic, p = 0.0229*), DR Midline vs Lateral 1 cluster (one-way ANOVA followed by Tukey multiple comparisons of means, F(2,219) = 6.022, p = 0.0028**; Chronic vs Acute, p = 0.0466*), and the whole dataset (no difference). Control n = 3, acute n=3, chronic n=3 mice per group. Dots represent individual spots in the ST dataset, p-values p < 0.05 = *, p < 0.01 = **, p < 0.001 = ***.

**Figure S5.**
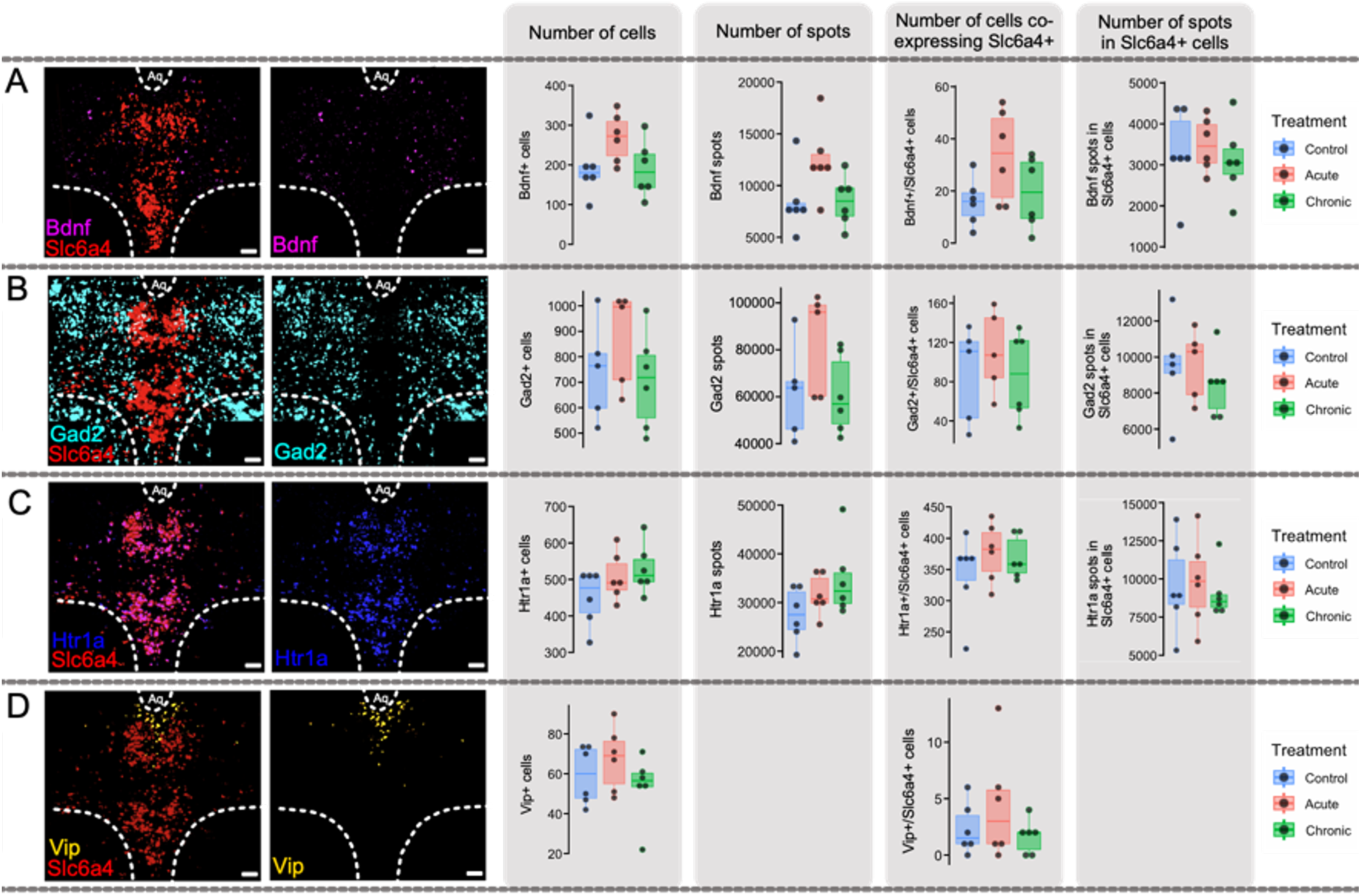
Gene expression of Bdnf, Gad2, Htr1a and Vip in the DR across SSRI treatment. **A.** Representative images of in-situ hybridization of *Bdnf* and *Slc6a4* co-expression and *Bdnf* in the DR. Box plots comparing treatment groups: sum of *Bdnf+* cells per section, sum of *Bdnf* spots (transcripts) per section, sum of cells co-expressing *Bdnf* and *Slc6a4* per section, and sum of *Bdnf* spots in *Slc6a4* cells. **B.** Representative images of in-situ hybridization of *Gad2* and *Slc6a4* co-expression and *Gad2* in the DR. Box plots comparing treatment groups: sum of *Gad2+* cells per section, sum of *Gad2* spots (transcripts) per section, sum of cells co-expressing *Gad2* & *Slc6a4* per section and sum of *Gad2* spots in *Slc6a4*+ cells. **C.** Representative images of in-situ hybridization of *Htr1a* and *Slc6a4* co-expression and *Htr1a* in the DR. Box plots comparing treatment groups of sum of *Htr1a+* cells per section, sum of *Htr1a* spots per slice, sum of cells co-expressing *Htr1a* & *Slc6a4* per and sum of *Htr1a* spots in *Slc6a4* + cells. **D.** Representative images of in-situ hybridization of *Vip* and *Slc6a4* co-expression and *Vip* in the DR. Box plots comparing treatment groups of sum of *Vip+* cells per section and sum of cells co-expressing *Vip* and *Slc6a4* per slice. Scale bar = 100μm, control n = 5-6, acute n=5-6, chronic n=6 mice per group.

**Figure S6.**
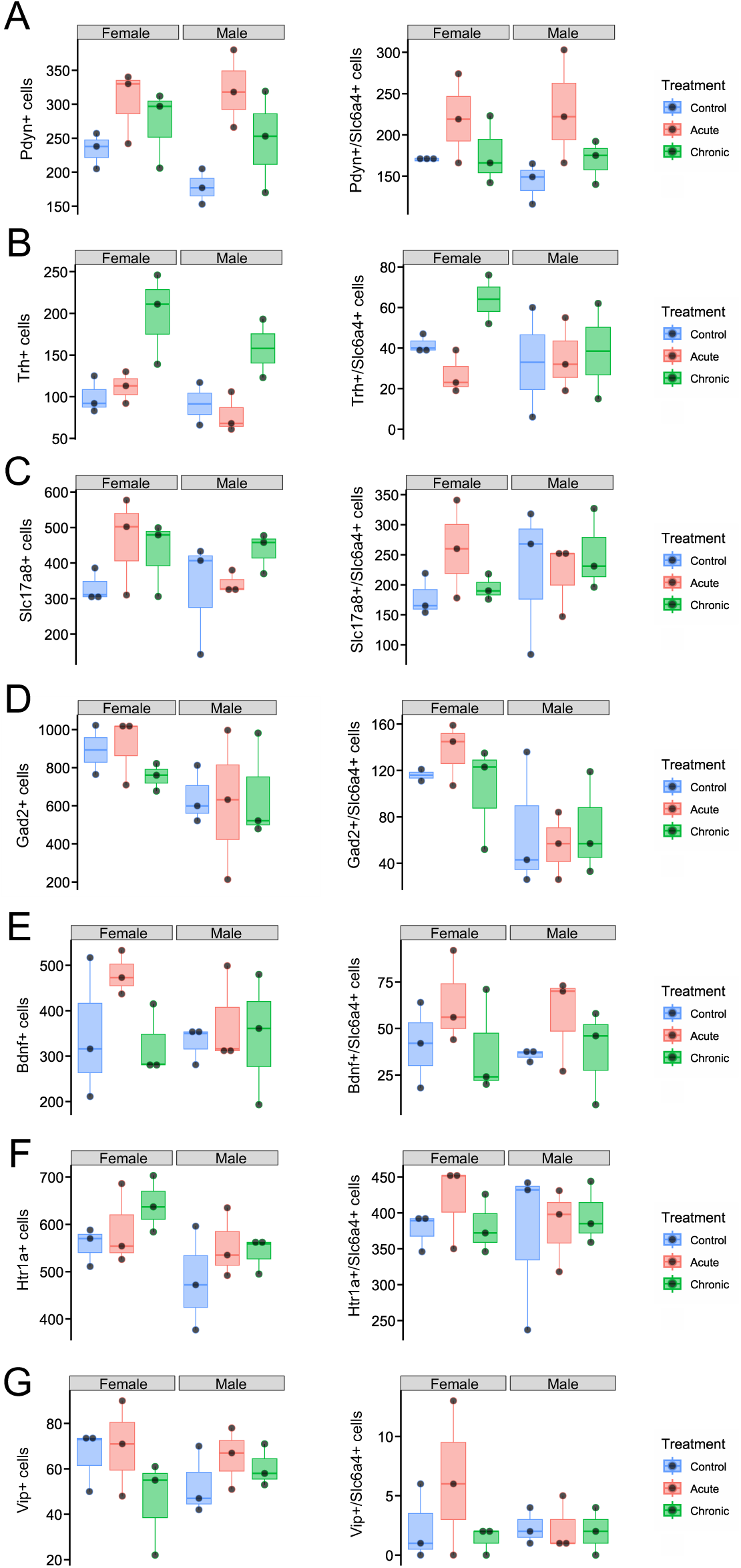
Lack of sex-specific modulation of gene expression in DR by SSRI treatment. No sex effects are observed due to acute nor chronic SSRI treatment in the expression of *Pdyn*, *Trh*, *Slc17a8*, *Gad2*, *Bdnf*, *Htr1a* and *Vip*. **A.** Box plots comparing treatment groups divided by sex: sum of *Pdyn+* cells per section, sum of cells co-expressing *Pdyn* and *Slc6a4* per section. **B.** Box plots comparing treatment groups divided by sex: sum of *Trh+* cells per section, sum of cells co-expressing *Trh* and *Slc6a4* per section. Control n = 5, acute n=6, chronic n=5/n=4 mice per group. **C.** Box plots comparing treatment groups divided by sex: sum of *Slc17a8+* cells per section, sum of cells co-expressing *Slc17a8* and *Slc6a4* per section. **D.** Box plots comparing treatment groups divided by sex: sum of *Gad2+* cells per section, sum of cells co-expressing *Gad2* and *Slc6a4* per section. Control n = 5, acute n=6, chronic n=6 mice per group. **E.** Box plots comparing treatment groups divided by sex: sum of *Bdnf+* cells per section, sum of cells co-expressing *Bdnf* and *Slc6a4* per section. **F.** Box plots comparing treatment groups divided by sex: sum of *Htr1a+* cells per section, sum of cells co-expressing *Htr1a* and *Slc6a4* per section. **G.** Box plots comparing treatment groups divided by sex: sum of *Vip+* cells per section, sum of cells co-expressing *Vip* and *Slc6a4* per section. Two-way ANOVA using Sidak multiple comparisons. Unless otherwise specified control n = 6, acute n=6, chronic n=6 mice per group.

